# Resonant Recognition Model analysis of cGAS: conserved spectral frequencies and a divergent naked mole-rat phase signature

**DOI:** 10.64898/2026.10.30.762874

**Authors:** Jose Luis Hernandez-Caceres, Mayrim Vega-Hernandez, Jorge Iglesias-Fuster, Ana Dorohova, Iván Padrón-Díaz

## Abstract

**Background:** Cyclic GMP–AMP synthase (cGAS) is a cytosolic DNA sensor with additional noncanonical nuclear functions, including suppression of homologous recombination (HR) in humans. In the naked mole-rat (NMR), cGAS instead promotes HR through a FANCI–RAD50-dependent mechanism, a functional reversal attributed to four C-terminal amino acid substitutions. Whether such functional divergence is reflected in primary-sequence-derived spectral features is unknown.

**Methods:** We applied the Resonant Recognition Model (RRM) to cGAS sequences from fourteen mammals and to FANCI sequences from eight species. Sequences were converted to numerical series using EIIP values and analyzed by discrete Fourier transform; consensus spectra were generated by point-wise multiplication of individual power spectra. Phase differences between cGAS and FANCI were computed at frequencies of interest. Spectral amplitudes at the dominant shared frequency were correlated with body mass, lifespan, and lifespan-to-body-mass ratio. RRM frequencies were converted to electromagnetic wavelengths using the Cosic formulation (λ = 201/f_RRM).

**Results:** The cGAS consensus spectrum contained four prominent peaks (f = 0.0392, 0.0765, 0.2539, 0.4229), with 0.2539 dominant. Three FANCI peaks (0.04138, 0.2479, 0.4332) were close to cGAS peaks, and the combined cGAS+FANCI consensus retained peaks at 0.2539 and 0.04077. Spectral amplitude at f = 0.2539 correlated with body mass (r = 0.647, p < 0.05) and with lifespan-to-body-mass ratio (r = −0.677, p < 0.01); the NMR showed the lowest amplitude. Human cGAS–FANCI phase difference was 3.1297 rad, 0.0118 rad from opposition, satisfying the RRM interaction criterion; the NMR cGAS–FANCI pair gave 1.7319 rad, approximately π/2, which does not satisfy the criterion. Conversion of the dominant frequency yielded a predicted wavelength of 792 nm (378.5 THz), in the near-infrared range. This prediction is testable in transparent vertebrate models: zebrafish respond to NIR light, and red-light pre-treatment has been reported to improve survival after lethal gamma irradiation in zebrafish embryos.

**Conclusions:** RRM identifies conserved spectral features in cGAS consistent with a cGAS–FANCI relationship, and amplitude at the dominant frequency varies with life-history traits. The phase analysis does not support a direct interaction signature for NMR cGAS–FANCI, a negative result consistent with a chromatin-mediated mechanism. The cross-species phase comparisons suggest that NMR FANCI, not only NMR cGAS, may contribute to the distinctive interaction mode. The predicted 792 nm wavelength provides a concrete, experimentally accessible parameter for testing whether RRM-derived frequencies can modulate radiation resistance in a transparent vertebrate model. These results are hypothesis-generating and require experimental validation.

## Introduction

The cyclic GMP–AMP synthase (cGAS) is best known as the cytosolic sensor of double-stranded DNA that synthesizes 2⍰3⍰-cGAMP and activates STING to induce type I interferon and inflammatory responses (Sun et al., 2013; Wu et al., 2013). This canonical axis is central to innate immunity against DNA viruses and intracellular bacteria. Over the past decade, however, cGAS has been found in the nucleus, at the plasma membrane, and in mitochondria, where it performs functions that are partly or wholly independent of interferon induction (Hopfner, et al., 2020; Qiu, et al., 2023, Bai et al., 2022).These noncanonical roles include suppression of DNA repair through chromatin compaction (Liu et al., 2018), membrane-associated self/non-self discrimination, and regulation of aging (Lu et al., 2024). Crucially, these activities are not uniformly catalysis-independent: nuclear DNA-repair suppression is largely catalysis-independent, whereas some aging-related phenotypes—such as ovarian aging driven by mitochondrial DNA leakage—require cGAMP production and STING signaling (Jiang et al., 2018).

The evolutionary history of cGAS reframes it as an ancient enzymatic scaffold rather than a vertebrate invention (Kranzusch et al. ,2015). The cGAS–STING cassette is deep-rooted, and STING’s capacity to bind cyclic dinucleotides predates type I interferon signaling (Margolis et al., 2017); invertebrate cGAS-like receptors (cGLRs) produce diverse cyclic dinucleotides in organisms lacking interferon genes, and the broader CD-NTase family has prokaryotic origins in antiphage defense (Kranzusch et al., 2015; Slavik & Kranzusch, 2023). Vertebrate-specific adaptations subsequently layered interferon induction and nuclear regulatory functions onto this conserved core (Morehouse et al., 2020).

The most striking illustration of cGAS functional divergence comes from the naked mole-rat (NMR), an exceptionally long-lived rodent (maximum lifespan ∼40 years) (Buffenstein and Jarvis, 2002). In human and mouse cells, nuclear cGAS is recruited to double-strand breaks, interacts with PARP1 via poly(ADP-ribose), and disrupts the PARP1–Timeless complex required for homologous recombination (HR), thereby suppressing accurate repair (Liu et al., 2018). NMR cGAS does the opposite: it does not engage the PARP1–Timeless axis, but instead promotes HR through enhanced FANCI–RAD50 interaction, and it does so independently of catalytic activity (Chen et al., 2025). This functional reversal is attributed to four amino acid substitutions in the C-terminal domain (S463D, E511K, Y527L, T530K), which reduce TRIM41-mediated K48-linked ubiquitination and P97 interaction after DNA damage, allowing prolonged chromatin retention that facilitates FANCI–RAD50 complex formation and RAD50 recruitment to damage sites.

These observations raise a question that sequence-level comparative analysis is well positioned to address: are the divergent functions of cGAS encoded as distinguishable features within the primary structure? The Resonant Recognition Model (RRM) offers one approach. RRM represents a protein sequence as a numerical series by assigning each residue a physical parameter—most commonly the electron–ion interaction potential (EIIP)—and applies discrete Fourier transform and cross-spectral analysis to identify frequency components shared among sequences with common biological function (Cosic, 1994). RRM has been used to locate functional “hot spots” in myoglobin, hemoglobin, and lysozyme, to predict mutational effects, and to design bioactive peptides (**Cosic et al, 1991; Krapez et al, 2025; Mishra et al., 2023)**.

Here we apply RRM to cGAS. We first derive consensus spectra from fourteen mammalian cGAS sequences and identify characteristic frequencies, then extend the analysis to FANCI proteins in order to test whether the two proteins share spectral features consistent with a functional relationship. Finally, we examine whether spectral amplitude at the dominant shared frequency varies across species in a manner correlated with body mass and lifespan, and we compare human and NMR cGAS directly to ask whether the four-substitution difference is detectable at the spectral level. We emphasize throughout that RRM is a heuristic predictor, and we frame its outputs as hypotheses requiring experimental validation.

## Methods

Sequences matching to cGAS proteins were downloaded from UniProt and corresponded to the following species:

1. A0A7N5P7A8 *Ailuropoda melanoleuca* (Giant panda, 517 AA)
2. Q8N884 *Homo sapiens* (Human, 522 AA)
3. A0A2U3XB78 *Leptonychotes weddellii* (Weddell seal, 522 AA)
4. A0A8C6C7B0 *Monodon monoceros* (Narwhal, 513 AA)
5. M3XZ01 *Mustela putorius furo* (Ferret, 526 AA)
6. A0A341ADK5 *Neophocaena asiaeorientalis* (Narrow-ridged finless porpoise, 513 AA)
7. A0AAX6RS70 *Heterocephalus glaber* (Naked mole-rat, 554 AA)
8. A0A8B7A7K3 *Orycteropus afer* (Aardvark, 523 AA)
9. A0A8C9CKN5 *Phocoena sinus* (Vaquita, 513 AA)
10. A0A671FHV0 *Rhinolophus ferrumequinum* (Greater horseshoe bat, 519 AA)
11. A0A2Y9DUW9 *Trichechus manatus* (American manatee, 523 AA)
12. A0A384BLH1 *Ursus maritimus* (Polar bear, 522 AA)
13. A0A3Q7SM56 *Vulpes vulpes* (Red fox, 521 AA)
14. A0A6J2DM97 *Zalophus californianus* (California sea lion, 522 AA)

Additionally, eight Fanconi anemia complementation group I (FANCI) amino acid sequences were downloaded, corresponding to the following species: naked mole rat, human, mouse, rat, chicken, *Arabidopsis thaliana*, panda, and wombat.

Sequences were converted into numerical series as per EIIP, in accordance with the method proposed by Cosic (1994) (Table 1). A MatLab script was produced for this purpose.

**Table 1.** EIIP values assigned to each amino acid. EIIP values assigned to Ile and as 0.0000 are by convention in the Cosic (1994) parameter set. These are not missing values.

| Amino acid | symbol1 | symbol2 | EIIP value |
|---|---|---|---|
| Alanine | Ala | A | 0.0373 |
| Cysteine | Cys | C | 0.0829 |
| Aspartic acid | Asp | D | 0.1263 |
| Glutamic acid | Glu | E | 0.0057 |
| Phenylalanine | Phe | F | 0.0946 |
| Glycine | Gly | G | 0.005 |
| Histidine | His | H | 0.0242 |
| Isoleucine | Ile | I | 0 |
| Lysine | Lys | K | 0.0371 |
| Leucine | Leu | L | 0 |
| Methionine | Met | M | 0.0823 |
| Asparagine | Asn | N | 0.0036 |
| Proline | Pro | P | 0.0198 |
| Glutamine | Gln | Q | 0.0761 |
| Arginine | Arg | R | 0.0959 |
| Serine | Ser | S | 0.0829 |
| Threonine | Thr | T | 0.0941 |
| Valine | Val | V | 0.0058 |
| Tryptophan | Trp | W | 0.0548 |
| Tyrosine | Tyr | Y | 0.0516 |

From numerical EIIP sequences, power spectra were obtained using MATLAB’s “pspectrum” command, and the consensus spectrum was generated by point-wise multiplication of the individual power spectra. The first 50 spectral values were discarded to mitigate the effect of the DC component. As result, spectral contributions below f=0.006 are not considered.

Consensus spectra were obtained for fourteen cGAS sequences, for eight FANCI sequences as well as for the total of fourteen cGAS sequences plus eight FANCI sequences

For phase analysis, in order to compute the phase difference between two putatively interacting proteins (the key RRM interaction criterion), complex Fourier sequences were obtained from individual EIIP sequences and phase differences were obtained for RRM frequencies of interest (Cosic, 1994). We focused on interactions between human and NMR cGAS sequences and FANCI sequences for NMR we tested the FANCI sequence G5CA91 · G5CA91_HETGA from Uniprot. We also tested the sequence <u>XM_004852819.1</u> from http://www.naked-mole-rat.org/annotations/details/XM_004852762.1/ and obtained identical results.

## Consensus spectra

The consensus spectrum from all fourteen cGAS sequences is represented in Figure 1. Four spectral peaks appeared at frequencies 0.0392(S/N=57.70), 0.0765(S/N=112.29), 0.2539(S/N=117.83), and 0.4229(S/N=15.48), with 0.2539 being the dominant peak.

**Figure 1.**
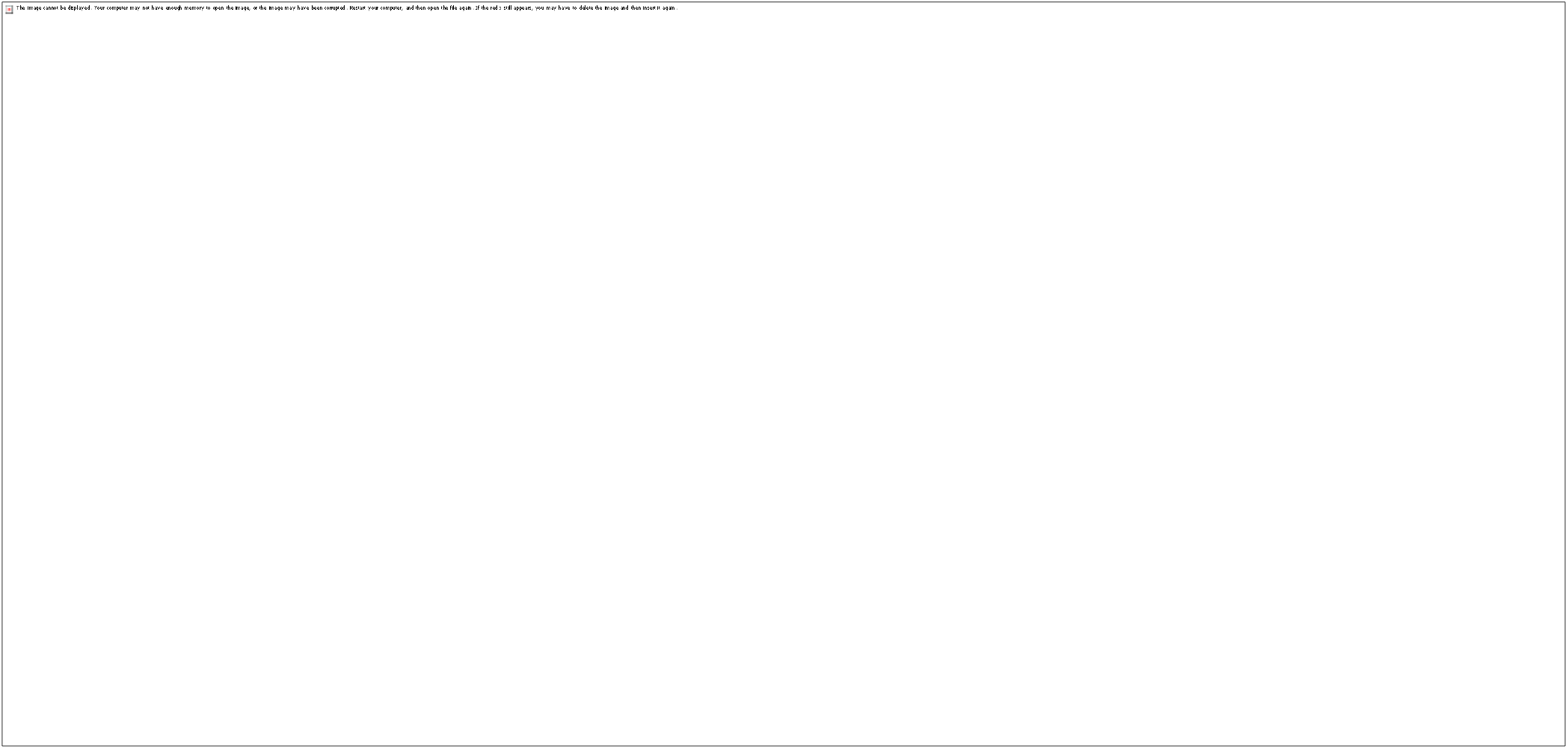
RRM consensus spectrum obtained from the fourteen cGAS sequences included in the study.

The consensus spectrum for the eight FANCI EIIP sequences is represented in Figure 2. Compared to cGAS, FANCI consensus spectrum exhibits a larger number of peaks, perhaps due to the smaller number of sequences. The first, prominent peak appears at f_RRM_=0.4408 (S/N=128.72). As documented by Cosic et al, this peak is close to the RRM frequency characteristic for microtubules(Cosic et al., 2014) Microtubules contribute to genome stability not only through mitotic chromosome segregation but also by facilitating DNA double-strand break repair (Ma et al, 2026). The γ-tubulin is recruited to damage sites and is required for efficient NHEJ and HR; its inhibition sensitizes cells to DNA-damaging agents, linking microtubule function to cellular radiation sensitivity (Solomon et al, 2025). More recently, KIF2A was shown to drive microtubule-dependent nuclear envelope invagination that creates a stable NHEJ environment and enhances 53BP1 recruitment at damage sites (Ma et al, 2026. In parallel, age-related decay of the neuronal microtubule cytoskeleton precedes other aging hallmarks, and genetic stabilization of microtubules slows neuronal aging in Drosophila (Okenve-Ramos et al, 2024). Together, these findings suggest that microtubule integrity supports the repair capacity required to survive genotoxic stress, and that its age-related decline may contribute to genome instability during aging (Rico et al, 2023).

**Figure 2.**
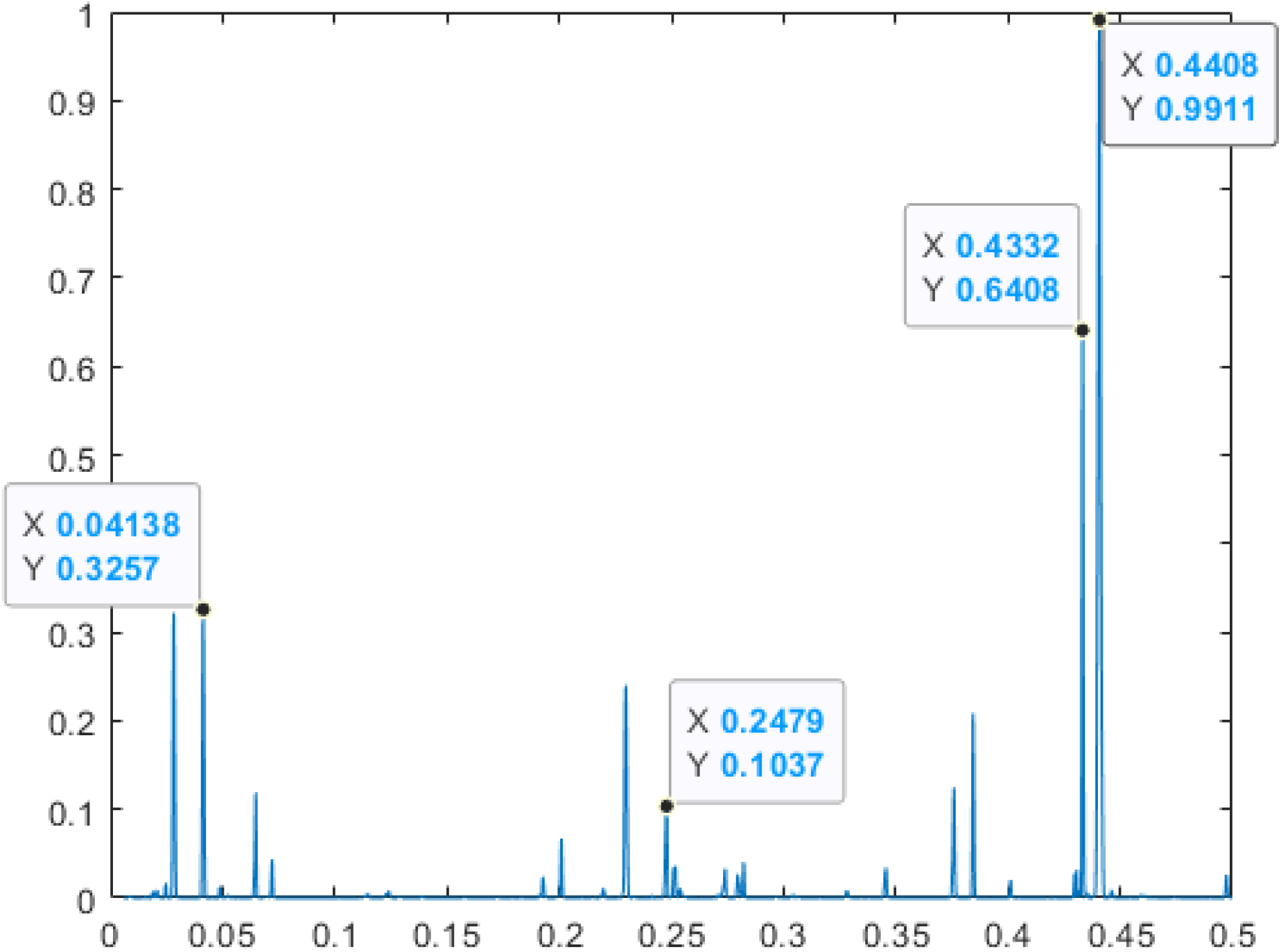
RRM consensus spectrum obtained from the eight FANCI sequences included in the study.

Noteworthy, three peaks in the FANCI consensus spectrum appear very close to three of the four peaks found in cGAS spectrum, apparently suggesting interaction between both proteins. These peaks are at the RRM frequencies 0.04138 (S/N= 42.30), 0.2479 (S/N= 13.45), and 0.4332 (S/N=83.23).

To further study the possible relation between cGAS and FANCI, a consensus spectrum was obtained for all sequences.

The resulting consensus spectrum (figure 3) contains only two prominent peaks. The main peak at frequency (f=0.2539; S/N= 466.78) coincides with the main peak of cGAS consensus spectrum. The second peak (f=0.04077; S/N= 60.17) is coincident with the third peak of cGAS consensus spectrum.

**Figure 3.**
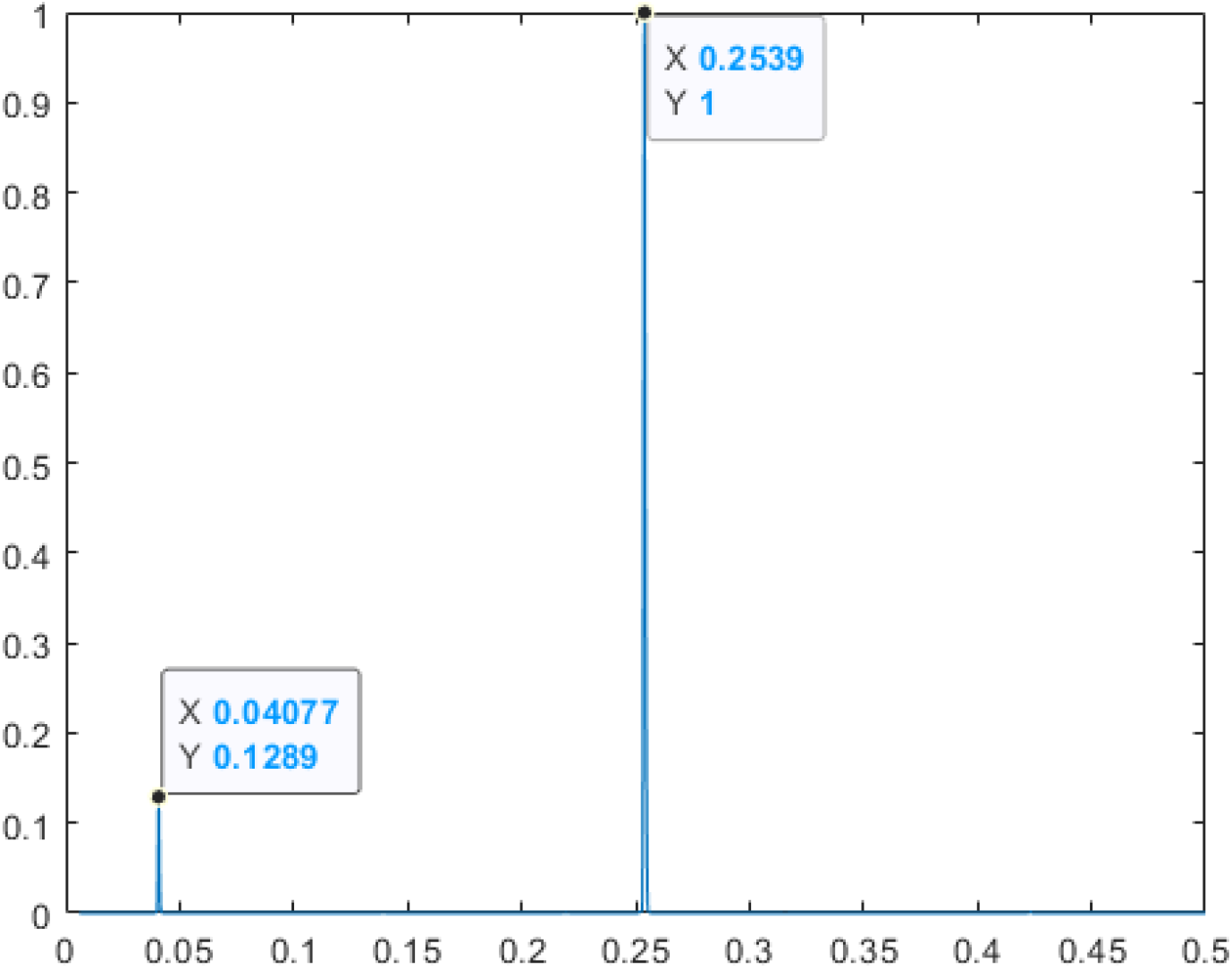
RRM consensus spectrum obtained from the fourteen cGAS sequences plus eight FANCI included in the study.

### 1. Spectral values at peak across species

#### Comparing spectral values at consensus cGAS-FANCI peak Across Species

The RRM frequency f=0.2539 corresponds, at the same time, to the main peak of the consensus spectrum of cGAS protein, as well as the main peak for the consensus spectrum of cGAS plus FANCI. This prominence suggests that this peak is associated to the interaction between cGAS and FANCI, which play a pivotal role in processes of regulation of DNA repair. Given the peculiar position of naked mole rat in these processes, it seems plausible to explore the amplitude of cGAS spectral peaks across species. Results are summarized in table2. As noticed, the shortest peak corresponds to naked mole rat. Spectral values are compared to lifespan, body mass and lifespan-to-body-mass ratio. Jackknife correlation analysis yielded significant correlations were found for spectral peak with body mass (r=0.647, p<0.05, CI [0.2427, 0.8594] and lor log-log dependence of lifespan-to-body-mass ratio (r=-0.677, p<0.05, CI [-1.4354, 0.2110], see figure 4).

**Table 2.**
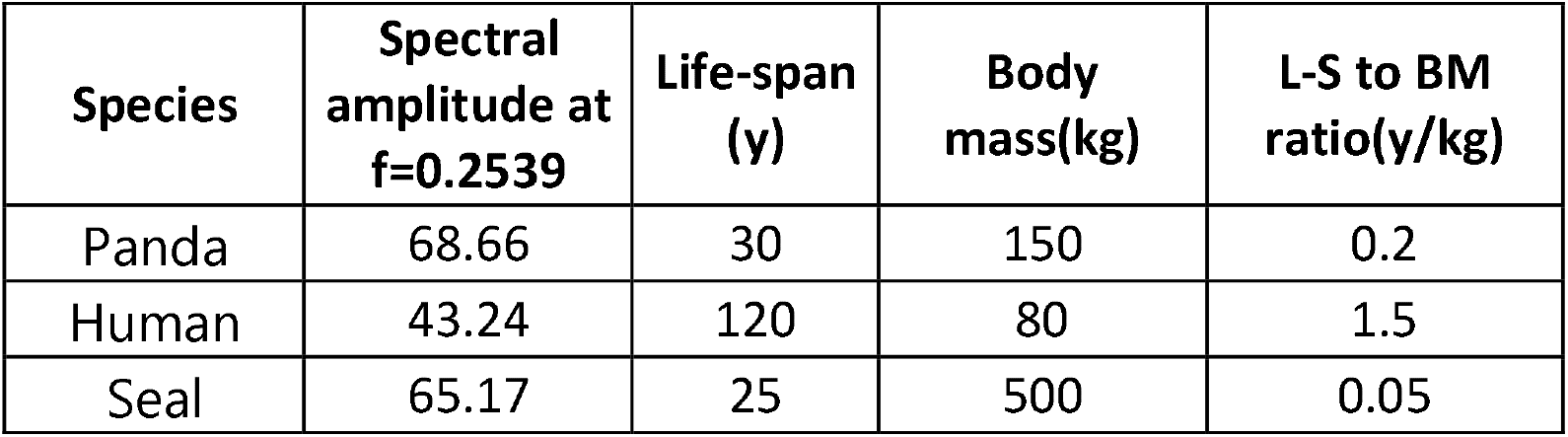
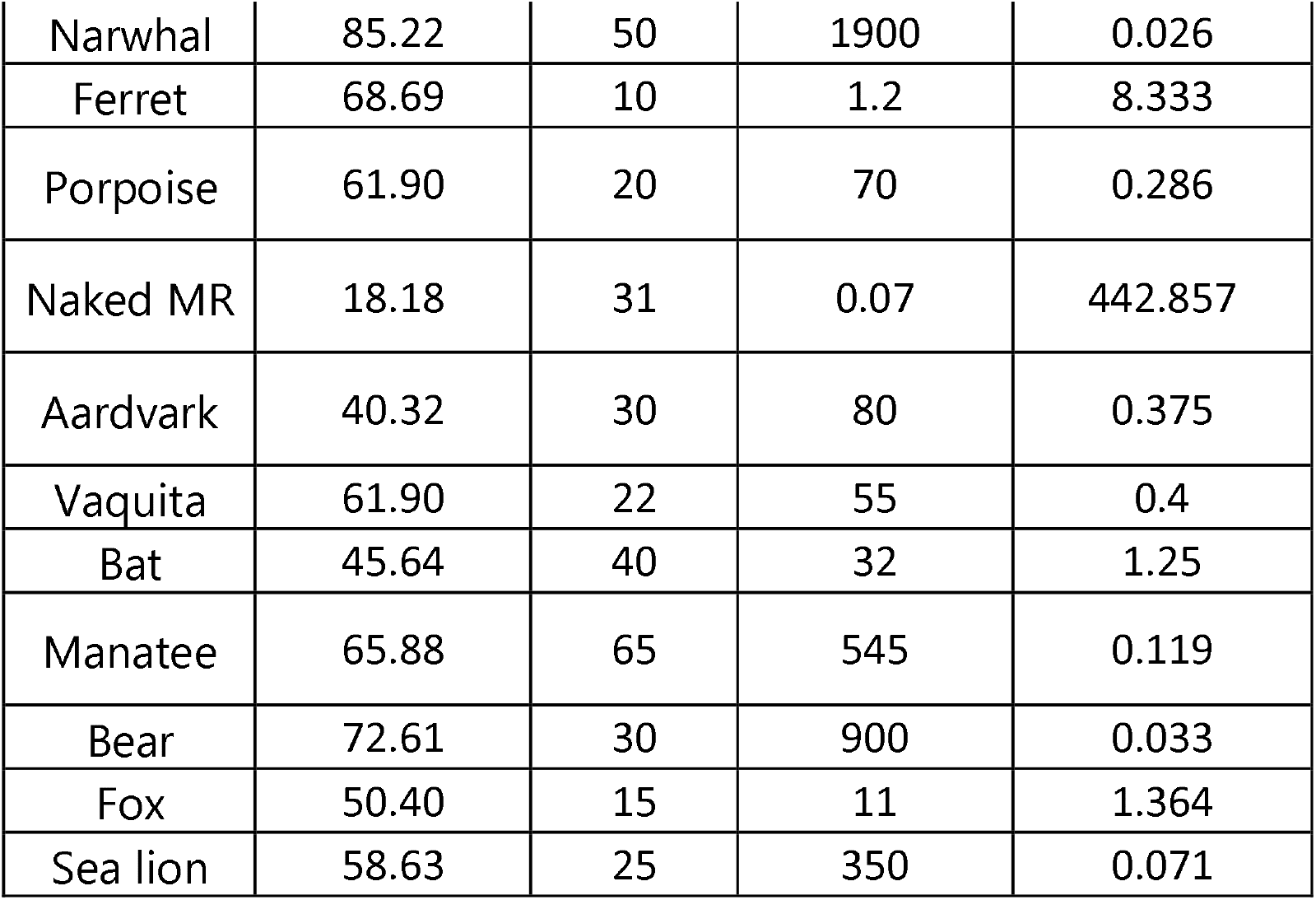
cGAS peak amplitudes at RRM frequency corresponding to cGAS-FANCI interaction (f=0.2539). Note that peak amplitude corresponding to NMR (highlighted in red) is the lowest, (twice smaller than the next in the list)

| **Species** | **Spectral amplitude at f=0.2539** | **Life-span (y)** | **Body mass(kg)** | **L-S to BM ratio(y/kg)** |
|---|---|---|---|---|
| Panda | 68.66 | 30 | 150 | 0.2 |
| Human | 43.24 | 120 | 80 | 1.5 |
| Seal | 65.17 | 25 | 500 | 0.05 |
| Narwhal | 85.22 | 50 | 1900 | 0.026 |
| Ferret | 68.69 | 10 | 1.2 | 8.333 |
| Porpoise | 61.90 | 20 | 70 | 0.286 |
| Naked MR | 18.18 | 31 | 0.07 | 442.857 |
| Aardvark | 40.32 | 30 | 80 | 0.375 |
| Vaquita | 61.90 | 22 | 55 | 0.4 |
| Bat | 45.64 | 40 | 32 | 1.25 |
| Manatee | 65.88 | 65 | 545 | 0.119 |
| Bear | 72.61 | 30 | 900 | 0.033 |
| Fox | 50.40 | 15 | 11 | 1.364 |
| Sea lion | 58.63 | 25 | 350 | 0.071 |

**Figure 4.**
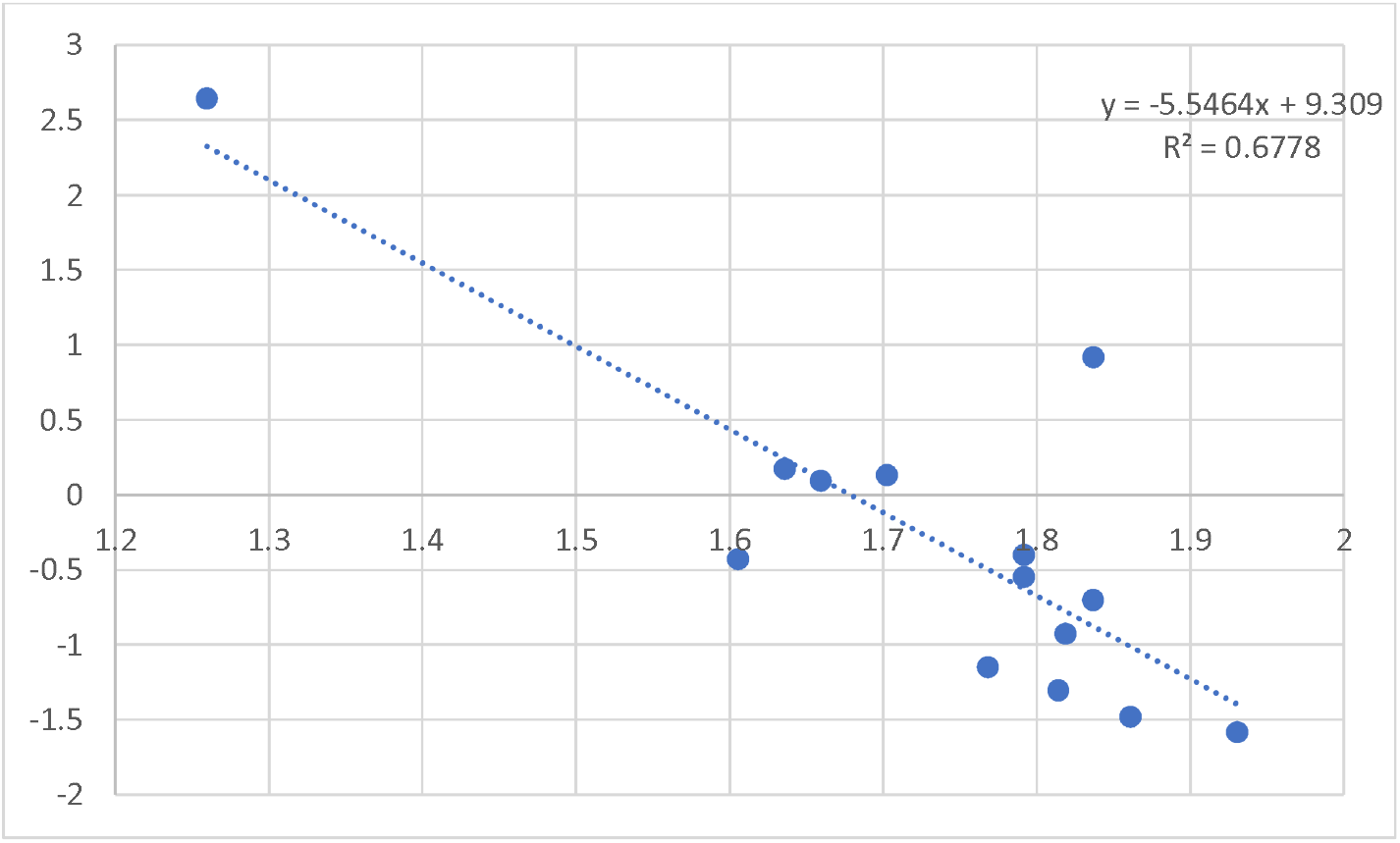
log-log plot for the relationship between amplitude of peak at RRM frequency 0.2539 and lifespan-to body mass ratio.

Even when pear amplitude is not a concluding criterion in RRM analysis, it can suggest some quantitative guidance. Thus, comparing RRM peak amplitudes with experimentally reported changes in HIV protease activity, Hernandez-Caceres et al found congruent results: those mutations increasing the RRM peak corresponded to increases in protease activity and vice-versa (Hernandez-Caceres et al, 2014).

### 1. Phase relationships

We searched for interaction between human cGAS and FANCI proteins at the frequency of the main peak. As it is known, in RRM analysis, phase relations are a strong criterion for interaction between proteins: in the RRM framework, the phase criterion for a putative direct interaction is phase difference near π (≈180°), i.e., opposite phase.

Results are summarized in table 3. As noticed, in humans the phase shift between cGAS and FANCI is almost opposite (3.1296 rad or 179.3°). Contrastingly, NMR cGAS and FANCI showed a phase difference of 99.2° very close to 90°.

**Table 3.** summary of phase difference analysis for interaction between cGAS and FANCI proteins from human and naked mole rat.

| **Pair** | **Phase difference (rad)** | **Distance to π (rad)** | **Interpretation.** |
|---|---|---|---|
| Human FANCI vs. Human cGAS | 3.1297 | -0.0118 | Only 0.0118 rad from π, satisfying Cosic's phase-opposition criterion for a putatively interacting pair between human cGAS and FANCI |
| NMR FANCI vs. NMR cGAS | 1.7319 | -1.4096 | Phase shift close to 90° (99.2°) is interpreted as a negative result for a direct cGAS–FANCI interaction at f = 0.2539 at NMR. This is consistent with the experimentally established indirect, chromatin-mediated mechanism. |
| Human FANCI vs. NMR cGAS | 2.9056 | -0.2360 | Suggest weakened, but still relevant, cGAS–FANCI interaction signature for cross species interaction. |
| NMR FANCI vs. Human cGAS | 1.5078 | -1.6338 | Suggest an absence of cross-species cGAS–FANCI interaction signature (86.4°, close to 90°) |

Noteworthy the fact that we obtained exactly the same values (phase difference ≈ 1.7319 rad, ∼90°) from two independent sources for the NMR FANCI sequence — the UniProt entry G5CA91 and the naked-mole-rat.org annotation XM_004852762.1 /XP_004852819.1 — means the result is robust and not an artifact of a single database entry. The two sequences are the same protein, so identical spectral output is exactly what you would expect. In the RRM framework, the phase criterion for a putative direct interaction is phase difference near π (≈180°), i.e., opposite phase, as the case for cGAS-FANCI interaction in Human. A value of π/2 (∼90°) sits halfway between “in phase” (0°) and “opposite phase” (180°). It is therefore not a signature of a direct resonant interaction at f = 0.2539. This is a negative result for the direct cGAS–FANCI interaction hypothesis under the standard RRM criterion.

At the same time, cross species comparisons suggest that cGAS from both Human and NMR are producing similar qualitative results. This, in our opinion, points to specific changes in the NMR FANCI protein. So far, we did not find studies comparing NMR FANCI proteins to other species, and this suggestion from RRM analysis could lead to relevant questions.

Noteworthy, the phase difference between human and NMR cGAS sequences is only 12.8 degrees, whereas the phase difference between human and NMR FANCI proteins is 265.7 degrees, close to -90 degrees. Though preliminary, these results, together with those from Table 3 suggest that the peculiar interaction between cGAS and FANCI is a result not only of changes in NMR cGAS protein.

#### Exploring Electromagnetic Modulation of Protein-Protein Interactions

Our analysis identifies f_RRM_ = 0.2539 as a major spectral contributor to cGAS–FANCI coordination. The Resonant Recognition Model (RRM) predicts an association between this numerical frequency and a physical electromagnetic wavelength. Using the Cosic formulation, the RRM frequency of 0.2539 corresponds to a wavelength of approximately 792 nm (378.5 THz). The conversion is defined by the empirical correlation established by Cosic (1994):

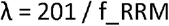

This relationship, with a slope factor K = 201, was derived from experimental correlations between computationally predicted frequencies and measured light absorption or irradiation effects in proteins. A wavelength of 792 nm falls in the near-infrared (NIR) region, just beyond the visible red limit, consistent with Cosic’s broader claim that RRM frequencies typically correspond to infrared and visible light ranges.

Direct experimental support for RRM-predicted photon emission comes from Karbowski et al. (2015), who measured a dominant emission peak at ∼385 nm from a bovine albumin solution, closely matching the ∼380 nm wavelength predicted by the RRM for this protein (Fig. 1B of that study).

Because the cGAS–FANCI interaction plays a role in genome stability and radioresistance, the RRM prediction that this interaction occurs at ∼792 nm suggests a testable hypothesis. It is technically feasible to expose a small, NIR-transparent animal model to monochromatic light at this wavelength.

Supporting this direction, researchers have used near-infrared wavelengths in the 800–810 nm range for various purposes in zebrafish. More directly, a 2024 conference paper investigated the radioprotective potential of red LED radiation using a zebrafish embryo model (dos Anjos Gusmão et al, 2024). The study found that pre-treatment with red light increased survival to over 50% after LD50 gamma radiation, suggesting that red LED could help reduce the acute effects of radiotherapy . While this study demonstrates the concept that light pre-treatment can modulate radiation resistance in zebrafish, it does not confirm that 792 nm would produce the same effect.

Zebrafish are sensitive to NIR light: research shows they respond behaviorally to wavelengths of 860 nm, and a broader comparative study found sensitivity to light at λ ≥ 750 nm in zebrafish. This confirms that 792 nm would be biologically relevant rather than passing through unnoticed. Together, these findings provide a rationale and methodological foundation for designing a study to test whether this specific frequency, derived from RRM analysis, can modulate radiation resistance in a transparent vertebrate model.

The search results confirm the authors, the LD50 dose (16.7 Gy), and the finding that pre-treatment increased survival to over 50% .

## Discussion

RRM analysis of fourteen mammalian cGAS sequences identified four consensus peaks, of which f = 0.2539 was dominant and was retained in the combined Casamance consensus. Three FANCI consensus peaks lay close to cGAS peaks, and phase analysis at f = 0.2539 placed the human cGAS–FANCI pair within 0.0118 rad of opposition, satisfying the RRM interaction criterion. The NMR cGAS–FANCI pair, by contrast, gave a phase difference of 1.7319 rad (∼π/2), which does not satisfy the criterion. Spectral amplitude at f = 0.2539 varied across species and correlated with body mass and lifespan-to-body-mass ratio, with the NMR showing the lowest amplitude. Cross-species phase pairings gave intermediate values, suggesting that the distinctive cGAS–FANCI relationship in NMR may involve changes in both proteins.

The four cGAS consensus peaks likely reflect distinct functional modules within the protein. The dominant peak at f = 0.2539 is shared with the cGAS+FANCI consensus and is therefore the strongest candidate for a frequency associated with the DNA-repair/nuclear interface. The second conserved feature, at f ≈ 0.040, appears in both the cGAS consensus and the combined consensus and may relate to the DNA-binding or catalytic core; this assignment is speculative and would require site-directed mutagenesis to test. The peaks at 0.0765 and 0.4229 are less clearly resolved and may reflect lineage-specific or function-specific contributions; the consensus approach necessarily obscures per-species variation, and individual spectra should be examined before these peaks are assigned functional meaning.

Chen et al. (2025) established experimentally that NMR cGAS promotes FANCI–RAD50 complex formation and RAD50 recruitment to damage sites. The RRM analysis reported here does not discover this relationship; its contribution is to ask whether the relationship leaves a detectable signature in the primary sequences. The answer is partial. The FANCI consensus spectrum shares three peaks with cGAS, and the combined consensus retains the dominant cGAS peak at f = 0.2539 with a high S/N (466.78), consistent with a shared spectral feature. However, the phase criterion—phase difference near π, as formalized by Cosic (1994)—is satisfied only for the human pair. The NMR pair sits near π/2, halfway between in-phase and opposite-phase, and therefore does not meet the criterion for a direct binary interaction at this frequency.

This negative result should not be read as evidence against a functional relationship between NMR cGAS and FANCI. The experimentally established mechanism is indirect: cGAS acts as a chromatin-retention platform that increases the local encounter probability of FANCI and RAD50, rather than binding FANCI through a direct interface. The RRM phase criterion was developed for globular protein–protein interfaces and would not be expected to detect a chromatin-scaffolded relationship. The negative phase result is therefore consistent with the known biology, and it illustrates a general caution for RRM applications: co-occurrence of a characteristic frequency in the consensus spectra of two proteins does not by itself imply a direct physical interaction.

The cross-species pairings in Table 3 provide additional information. Human FANCI paired with NMR cGAS gives a phase difference of 2.9056 rad (166.5°), 0.236 rad from opposition—a weaker but still opposition-side match. NMR FANCI paired with human cGAS gives 1.5078 rad (86.4°), which does not approach opposition. Taken together with the observation that the human and NMR cGAS sequences differ in phase by only 12.8° while the human and NMR FANCI sequences differ by 265.7° (equivalent to −94.3°), these results suggest that the distinctive cGAS–FANCI relationship in NMR is not attributable to cGAS alone. The FANCI protein may carry sequence features that contribute to the altered interaction mode. We are not aware of published comparative analyses of NMR FANCI, and this suggestion from RRM analysis could motivate such a study.

### Future directions

Four lines of work follow. First, experimental mutagenesis of RRM-predicted hot spots would test whether the consensus frequencies correspond to functionally important residues. Second, a windowed or pairwise spectral analysis restricted to the C-terminal regulatory region would isolate the effect of the four substitutions. Third, a phylogenetically informed analysis (e.g., phylogenetic generalized least squares) of amplitude versus life-history traits would account for shared ancestry. Fourth, extension of the analysis to cGLRs and prokaryotic CD-NTases would test whether the f = 0.2539 frequency predates interferon signaling. Finally, as suggested, zebra fish experiments using electromagnetic waves in the near infrared could suggest a new way of exploring radiation resistance mechanisms and possible modifications.

## Conclusion

RRM recovers conserved spectral features in cGAS and a shared signature with FANCI, and amplitude at the dominant frequency tracks life-history traits across mammals. The phase analysis, however, does not support a direct cGAS–FANCI interaction in the NMR, a negative result consistent with the chromatin-mediated mechanism established experimentally. The cross-species phase comparisons suggest that NMR FANCI, not only NMR cGAS, may contribute to the distinctive interaction mode. These findings are hypothesis-generating, and point to specific mutagenesis and zebrafish radiation resistance experiments.

